# A benchmark of Hi-C scaffolders using reference genomes and *de novo* assemblies

**DOI:** 10.1101/2022.04.20.488415

**Authors:** Aakash Sur, William Stafford Noble, Peter J. Myler

## Abstract

**Background:** Studying a new species using high-throughput sequencing requires a high-quality reference genome. However, assembling chromosome length sequences remains challenging. Recent advances in chromatin conformation capture (Hi-C) have provided a new approach to scaffolding genome assemblies, and the last ten years have seen a proliferation of such methods. However, to our knowledge no comprehensive benchmarking of Hi-C scaffolders has been conducted to date.

**Results:** Through a literature review we identified the most popular Hi-C scaffolders – Lachesis, HiRise, 3d-dna, SALSA, and AllHiC. We tested their ability to scaffold four well studied genomes – *S. cerevisiae, L. tarentolae, A. thaliana*, and *H. sapiens*. Scaffolders were tasked with both scaffolding fragmented versions of the reference genome as well as *de novo* assemblies derived from long read datasets. We found that all scaffolders can exceed 80% accuracy under ideal circumstances but that their performance quickly deteriorates under more challenging conditions. Surprisingly, many scaffolders also showed poor performance on the best assemblies, where contigs are near chromosome length. Overall, we found that HiRise and Lachesis offer the best performance on average across all conditions.

**Conclusions:** We compare the performance of five Hi-C scaffolders using multiple reference species under both ideal and real-life conditions, thereby illuminating their strengths and weaknesses.

## Introduction

Robust genome sequences are foundational to molecular biology, yet many reference genomes remain in the purgatory of the draft assembly. Producing complete chromosome-length sequences of organisms has long been the goal for genome assembly, but progressing from high-confidence contigs to scaffolded chromosomes has proven challenging. Recent experimental advances in interrogating the three-dimensional structure of genomes through chromatin conformation capture (Hi-C) has provided valuable new information to help solve the assembly problem [1]. Several groups have developed scaffolding algorithms that utilize Hi-C data to group, order, and orient contigs into completed genomes. Despite the proliferation of such methods, the accuracy of these methods has never been comprehensively benchmarked.

A search of the current literature revealed ten Hi-C scaffolding methods, of which five (Lachesis, HiRise, 3d-dna, SALSA, and AllHiC) have been used in a publication more than three times (Table 1). In each of their respective publications, the authors test their scaffolder by solving a genome assembly, but the species and task vary considerably among papers. Occasionally, a reference is artificially split into an arbitrary number of pieces and the scaffolder tasked with reassembling it [6,7]. In some cases assemblies of known genome are scaffolded [1,4,5,6,7], and other times unpublished genomes are solved [5,6,7].

**Table 1.** The Hi-C scaffolding tools identified in the literature search along with their publication date, citation count, and number of times they were used in a genome publication as of September 15, 2020. Although scaffolders published earlier tend to have more citations, their actual application count varies significantly. Our selection criteria required scaffolders to have three or more applications. *“Manual annotation” refers to studies that complete the genome by hand using the Hi-C map as a visual guide.

| Name | Publication Date | Citation Count | Application Count |
| --- | --- | --- | --- |
| Lachesis [1] | December 1, 2013 | 481 | 89 |
| Dna-triangulation [2] | December 1, 2013 | 138 | 0 |
| Graal [3] | December 17, 2014 | 108 | 1 |
| HiRise [4] | February 4, 2016 | 349 | 17 |
| 3d-dna [5] | April 7, 2017 | 297 | 40 |
| Salsa [6] | July 12, 2017 | 71 | 5 |
| ALLHiC [7] | August 5, 2019 | 20 | 4 |
| HiCAssembler [8] | October 10, 2019 | 8 | 0 |
| HiC-Hiker [9] | May 5, 2020 | 0 | 0 |
| Manual Annotation* | N/A | N/A | 5 |

Our primary objective in this study is to evaluate the performance of existing Hi-C scaffolders using a uniform set of tests and evaluation metrics. To this end, we evaluated the five most commonly used Hi-C scaffolders against four diverse eukaryotic species — *Saccharomyces cerevisiae, Leishmania tarentolae, Arabidopsis thaliana*, and *Homo sapiens*. Each of these genomes has unique characteristics, including widely varying genome size, base composition, rate of interchromosomal interaction, and even evolutionary differences in chromosome packing.

We tested the scaffolders with two different sets of contigs: split reference and *de novo* assemblies. The split assembly divides a reference genome into equal-sized pieces, presenting an artificially pristine test that is the best-case scenario for each scaffolder. Because real genome assembly is usually complicated by repeat ambiguity, haplotypes, and low complexity sequences, we created several *de novo* assemblies from long read datasets using the Canu assembler. [10] In both settings, scaffolders are tasked with joining a given set of contigs to produce scaffolds, which are then compared against the existing reference genome. We observed that scaffolders performed better on average on the split reference task than the *de novo* assembly task. Additionally, we found that the accuracy of the scaffolders changed with the species being tested, suggesting that sequence characteristics such as repeat content and heterozygosity levels may affect performance. On average, HiRise and Lachesis performed the best, with HiRise and Salsa working best on less fragmented assemblies, and HiRise, Lacheis, or AllHiC being better choices for more fragmented assemblies. Although scaffolders can perform well under ideal circumstances, our results suggest that existing Hi-C scaffolders are still expected to make mistakes, requiring manual correction before new reference genomes can be published.

## Methods

### Literature Search

To find all available Hi-C scaffolders, we conducted a literature search on PubMed for publications between January 1, 2010, to September 15, 2020. The search terms “(hi-c scaffolding) or (hi-c assembly) or (hi-c genome assembly) or (hic scaffolding) or (hic assembly) or (hic genome assembly)” yielded 370 results, of which 171 were ultimately deemed relevant (Supplementary Table 1). Ten scaffolding methods were identified, as well as the frequency with which they have been used to publish genomes. Methods with three or more cited applications were selected for benchmarking.

### Split reference and de novo assemblies

To generate the split assemblies, we partitioned the established reference genomes of *S. cerevisiae, L. tarentolae, A. thaliana*, and *H. sapiens* into equal sized pieces of 10kb, 50kb, 100kb, 500kb, and 1mb to create a total of sixteen split reference assemblies. To generate a diverse set of *de novo* assemblies for each of the four organisms, we collected a large repository of long-read data from NCBI’s SRA database, as well as our previously generated PacBio reads for *L. tarentolae* (see Supplementary Table 3). To normalize for different genome sizes, we down-sampled the number of reads to achieve a theoretical coverage of 10x, 20x, …, 100x for each reference genome. We ran the Canu assembler on each of these down-sampled datasets using the default parameters to create forty different *de novo* assemblies summarized in Supplementary Table 4. Canu was selected given its popularity as a long-read assembler as well as its robust support for clusters and scheduling frameworks such as Slurm.

### Hi-C Alignments

Hi-C scaffolders use the alignment of Hi-C reads against a genome assembly to optimally place contigs. We downloaded publicly available datasets of Hi-C reads from the SRA database for *S. cerevisiae, A. thaliana*, and *H. sapiens*, and used our previously generated Hi-C reads for *L. tarentolae* (Table 2). The Hi-C dataset for the human genome proved too large to easily work with so we opted to downsample that dataset to 100 reads per kilobase. All reads were aligned against *de novo* and split reference assemblies using BWA [11], and duplicate reads were filtered with samblaster [12]. The resulting alignment files were sorted with samtools [13] into either coordinate-sorted files or read-sorted files depending on scaffolder requirements. 3d-dna required the use of the Juicer pipeline [14], so reads were extracted from the filtered BAM alignments to be input to Juicer.

**Table 2.**
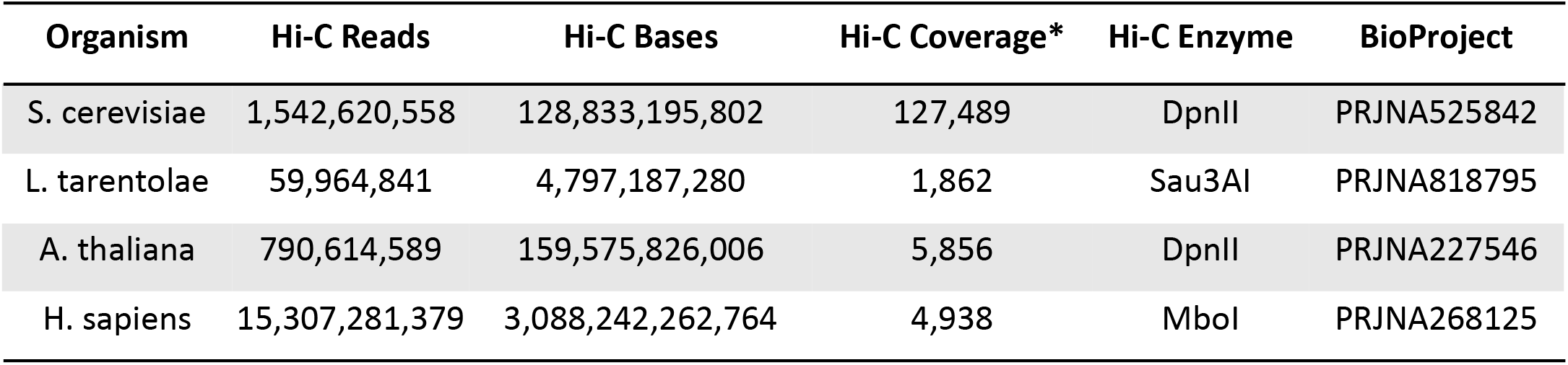
Overview of the Hi-C data collected for each of the four organisms. All the Hi-C enzymes cut at the same site, allowing the pipeline to remain consistent. The *H. sapiens* data proved to be computationally unwieldy, it was downsampled to 100 reads/kb. *Hi-C coverage is reported as reads per kilobase since the number of bases in a particular read do not contribute towards the contact count.

### Hi-C Scaffolding

Of the five scaffolders selected for benchmarking, two initially failed to build due to errors in the source code (Lachesis and HiRise), one required deprecated software dependencies (Salsa), and two installed as intended (3d-dna and AllHiC). To work with this diverse set of software tools and with an eye towards providing a community resource, we containerized the four methods with installation hurdles in Docker and have made them freely available (https://hub.docker.com/u/aakashsur). Scaffolders were then tasked with joining the contigs of a genome assembly using the alignment of Hi-C reads to that assembly. Lachesis and AllHiC require an expected number of chromosomes when running, representing a potential limitation of those methods. Additionally, HiRise was originally developed for use with in vitro chromatin proximity ligation reads (Chicago) rather than Hi-C data.

To determine how many Hi-C reads were required to effectively scaffold genomes, we selected a single assembly for each species and down-sampled the number of aligned and deduplicated Hi-C reads to 1, 50, 100, 500, and 1000 reads per kilobase. Since the difficulty of the scaffolding problem is often related to the fragmentation of the underlying genome assembly, we also wanted to determine how assembly quality affects scaffolding. This fragmentation is most often measured by the N50 of an assembly, which is the size at which contigs of equal or greater length cover half the assembly. In order to test the effects of different N50s, the number of Hi-C reads were normalized by downsampling each run to 100 aligned Hi-C reads per kilobase.

### Scaffolding Accuracy

Each scaffolder outputs a FASTA file where the appropriate joins have been made to the input contigs. To evaluate how closely this layout matches the optimal layout, we used Mummer 4 to map the assembly contigs to the known reference genome, and then compared them. [15] Using the python package Edison, we calculate the edit distance, overall accuracy, grouping accuracy, ordering accuracy, and orientation accuracy. [cite] The edit distance is the number of edits needed to alter a given set of scaffolds such that they most closely resemble a reference genome. This distance is calculated using the Double Cut and Join (DCJ) algorithm for genomic rearrangements, which guarantees finding the theoretical minimum edit distance [16]. The overall accuracy is obtained from the DCJ model by determining what fraction of sequence in the assembly has been correctly placed. The grouping accuracy measures how effectively a scaffolder is able to partition contigs into their associated chromosomes, and is computed using a length weighted Jaccard index between scaffolds and chromosomes. The ordering accuracy is the length-weighted frequency of finding a pair of adjacent contigs in the assembly that are also adjacent in the reference. Finally, the orientation accuracy is similar to the ordering accuracy, but also requires that adjacent contigs be correctly oriented relative to each other.

## Results

We benchmarked the five most utilized Hi-C scaffolders: Lachesis, HiRise, 3d-dna, SALSA, and AllHiC. To simulate ideal conditions, we challenged the scaffolders to reproduce the high-quality reference genomes of *S. cerevisiae, L. tarentolae, A. thaliana*, and *H. sapiens* that had been split into equal size pieces. In addition, to assess performance in a more realistic setting, we benchmarked each scaffolder using *de novo* assemblies which approximate the reference genomes but contain the ambiguities and complexities of real assemblies. (Table 3)

Several of the scaffolders proved challenging to install and run, so we have released patched versions as Docker containers at https://hub.docker.com/u/aakashsur. Despite these patches, there are inherent limitations to some of the software tools that lead to failed runs. Most commonly, Lachesis fails to finish if it is unable to build the specified number of chromosomes. Additionally, several scaffolders failed to complete the 10kb N50 split reference assembly of the human genome in the allotted 10 days — the maximum our cluster allows. In most cases, HiRise tends to have a run time that is an order of magnitude higher than the other scaffolders, and Lachesis consistently had the fastest time, even being able to scaffold human-sized genomes within an hour (Supplementary Table 9).

To determine the impact of Hi-C read coverage on scaffolding ability, we downsampled the number of Hi-C reads. As a baseline, we chose the 100kb split reference assembly to represent a plausible, modern assembly attempt. We found that the vast majority of the time, performance degrades as the amount of Hi-C data is reduced (Figure 1). Although we observe heterogeneity as to when this shift occurs, 50 reads/kilobase appears to be the point below which at least some of the scaffolders begin to perform worse. Lachesis and AllHiC appear to have both the smallest drop in performance suggesting that they are the most tolerant of less data. These results are broadly replicated in the *de novo* assembly setting as well (Supplementary Table 10).

**Figure 1.**
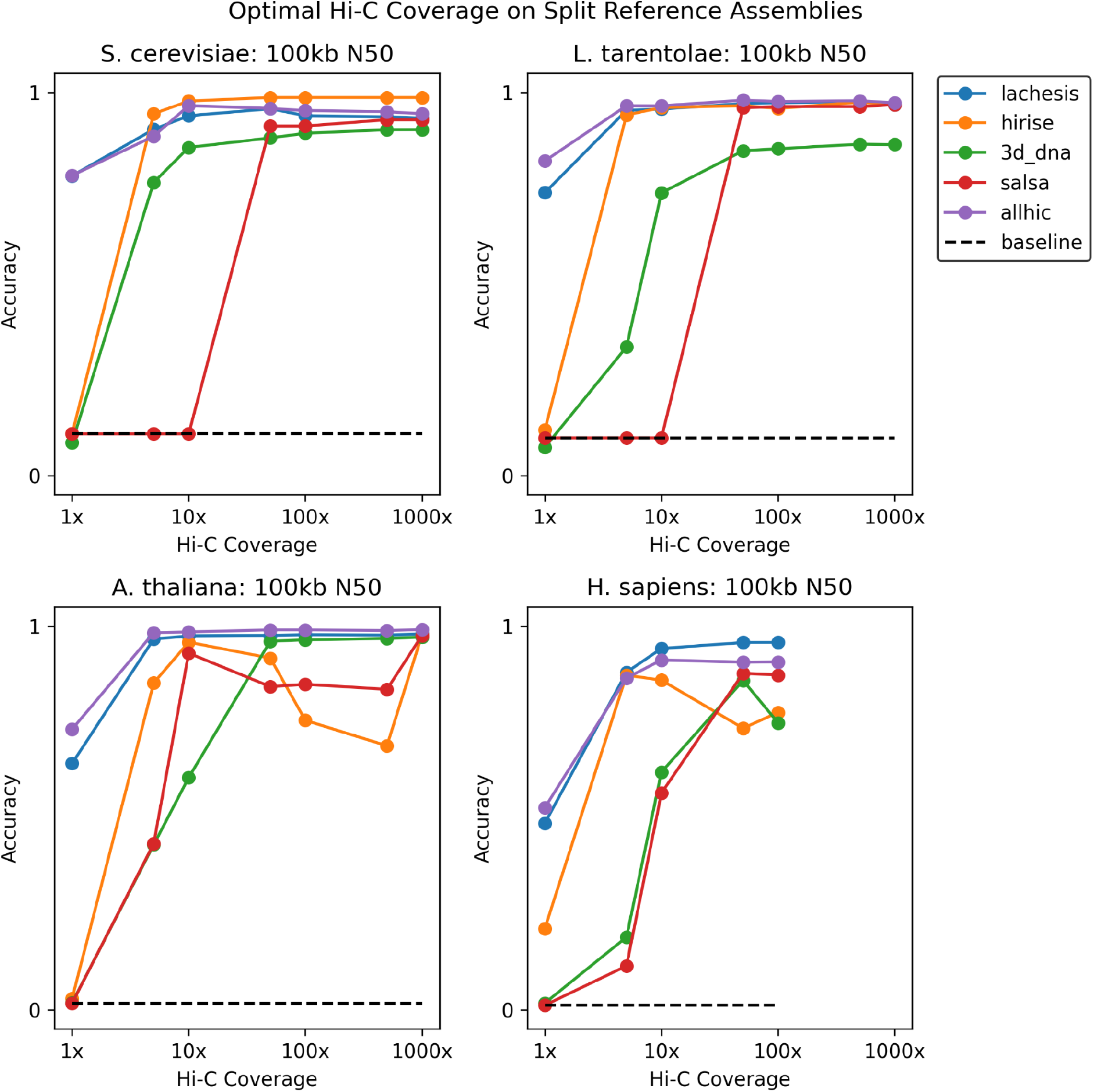
The effect of down-sampling Hi-C coverage on scaffolding accuracy. While using the 100kb N50 split reference assembly, reads were downsampled to target densities. Past 50 reads per kilobase, performance of all scaffolders tends to degrade, though some scaffolders are more resistant to decline than others.

To measure accuracy as a function of assembly size, we varied the N50 of the genome assemblies while keeping a constant 100 reads per kilobase of Hi-C data. For the split reference assemblies, we found that scaffolders can reach upwards of 80% accuracy in many cases, but occasionally perform much worse (Figure 2). Indeed, the best scaffolder for one species was not necessarily the best for another, making the calculus of ranking more challenging. Nevertheless, a consistent trend was lower performance on 10kb N50 assemblies, suggesting their high degree of fragmentation makes them difficult to scaffold. Though this might suggest that more contiguous assemblies ought to fare better, we were surprised to find that several of the scaffolders had dramatic decreases in performance with the highest N50s. In fact, on five occasions, scaffolders perform worse than the baseline of no scaffolding, indicating the errors they have created exceed the starting errors. For AllHiC, this decrease occurs because the method produces a single scaffold containing all the contigs (Supplementary Figure 12). Similarly, 3d-dna yielded consistently low grouping scores for these two species and also showed a decline in ordering accuracy at the high N50 range with similar problems in “over-scaffolding,” i.e., producing fewer scaffolds than there are chromosomes.

**Figure 2.**
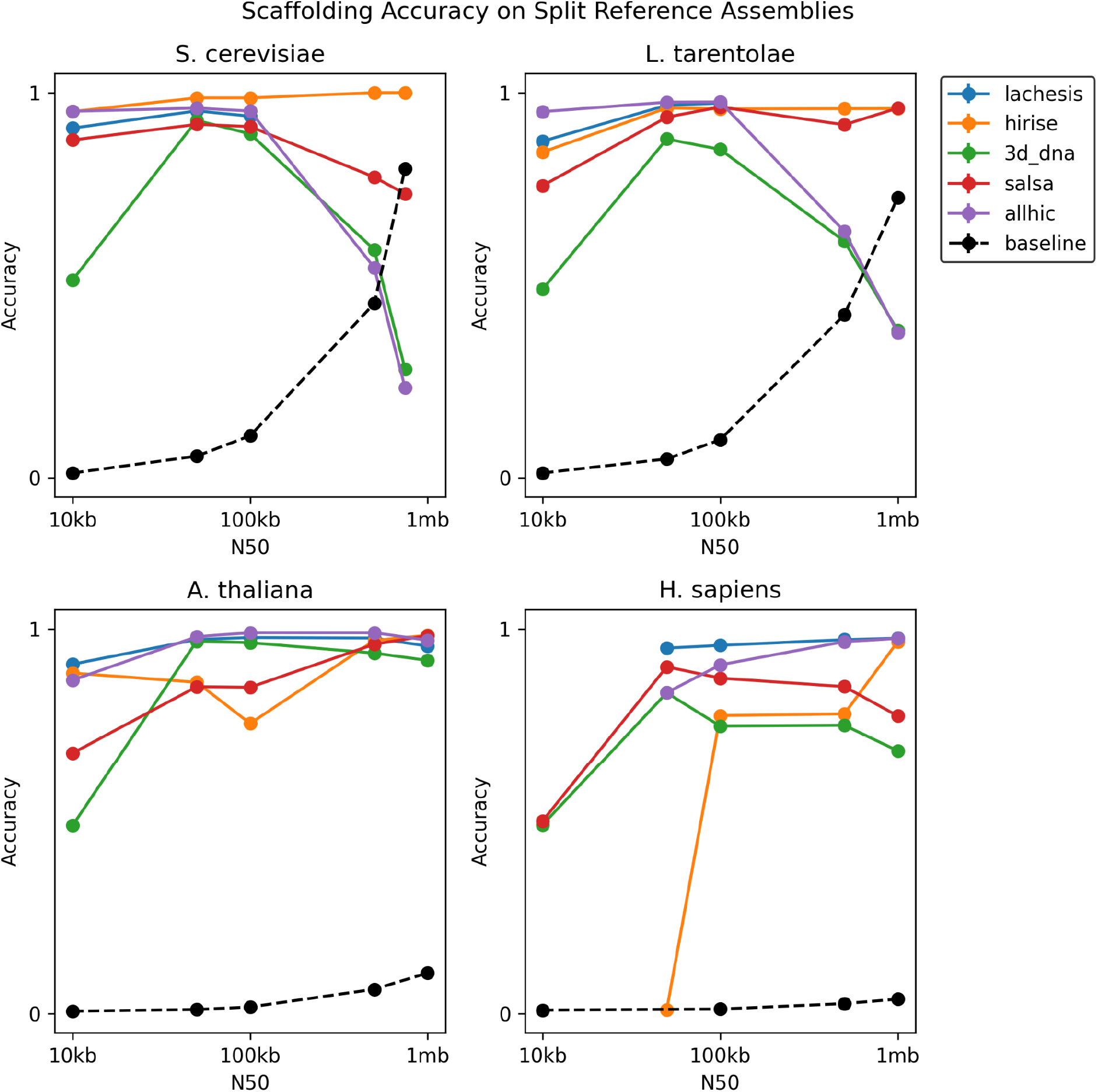
The accuracy of Hi-C scaffolders on four reference genomes. For each species, we created five different assemblies by splitting the reference in equal sized parts. High performance on a particular organism does not guarantee high performance on others. Performance decreases by 10kb N50 for all species, but also decreases at the high N50 range for *S. cerevisiae* and *L. tarentolae*.

For the *de novo* assemblies, significant variability in accuracy was observed across scaffolders and species, presumably due to the inherent complexities of the assembly process (Figure 3). As a general trend, the scaffolders tended to perform worse on *de novo* assemblies than they did on split references (Table 3). Particularly for *A. thaliana* and *H. sapiens*, accuracies are much lower and closer to baseline than in the split reference setting. Overall, we still see the trend where accuracy decreases at either extreme of the N50 spectrum, albeit in a less consistent fashion. Again, AllHiC and 3d-dna perform significantly worse on the more contiguous *S. cerevisiae* assemblies, similar to their trends on the split reference task.

**Figure 3.**
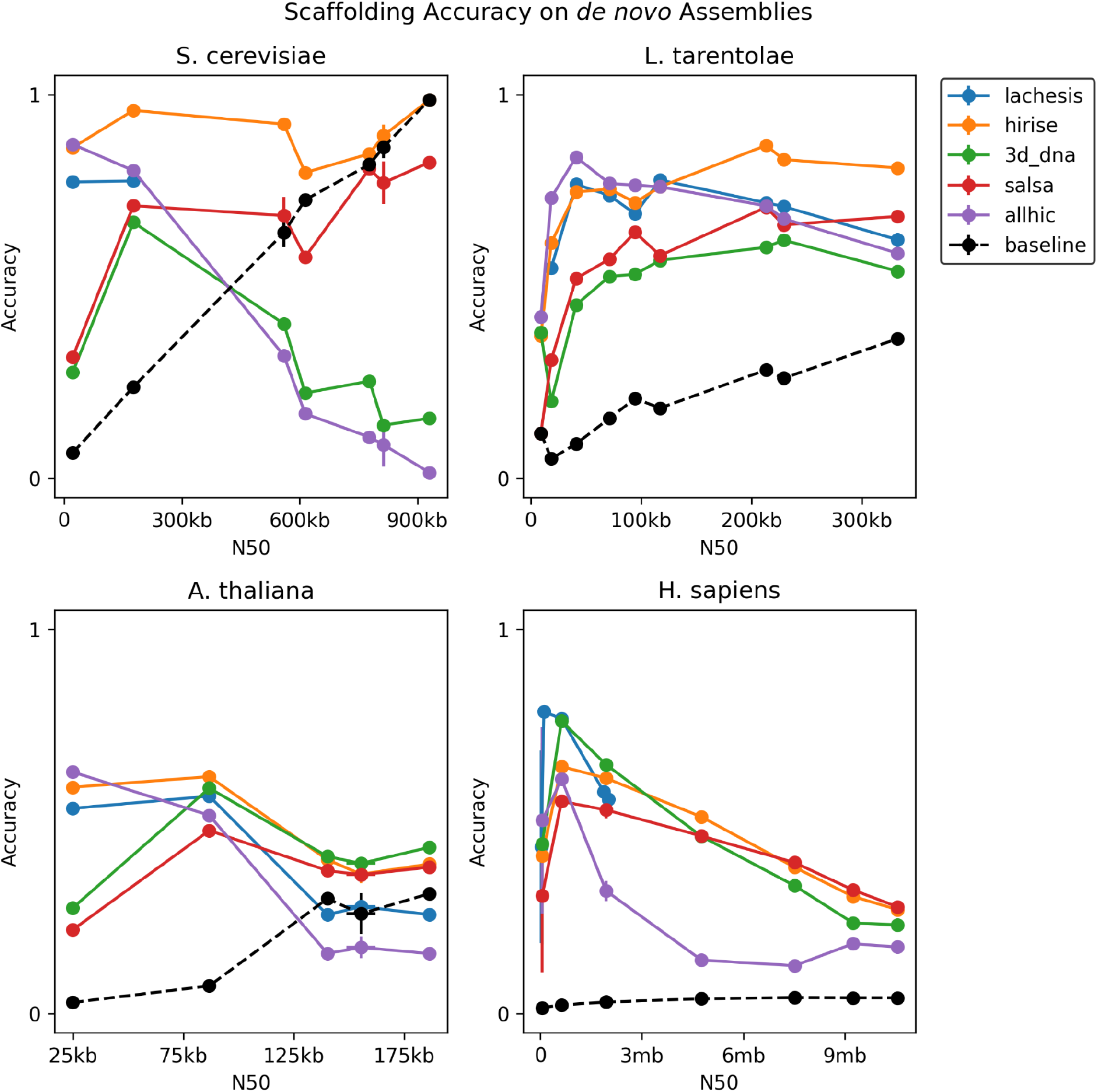
The accuracy of Hi-C scaffolders on the *de novo* assemblies of four species. The scaffolders exhibited significant variability across species, as well as an overall lower performance compared to the split reference reconstruction task. Nevertheless, a similar trend of poorer performance at either end of the N50 spectrum remains, with highly fragmented assemblies causing poorer performance, and highly contiguous assemblies causing a drop as well.

**Figure 4.**
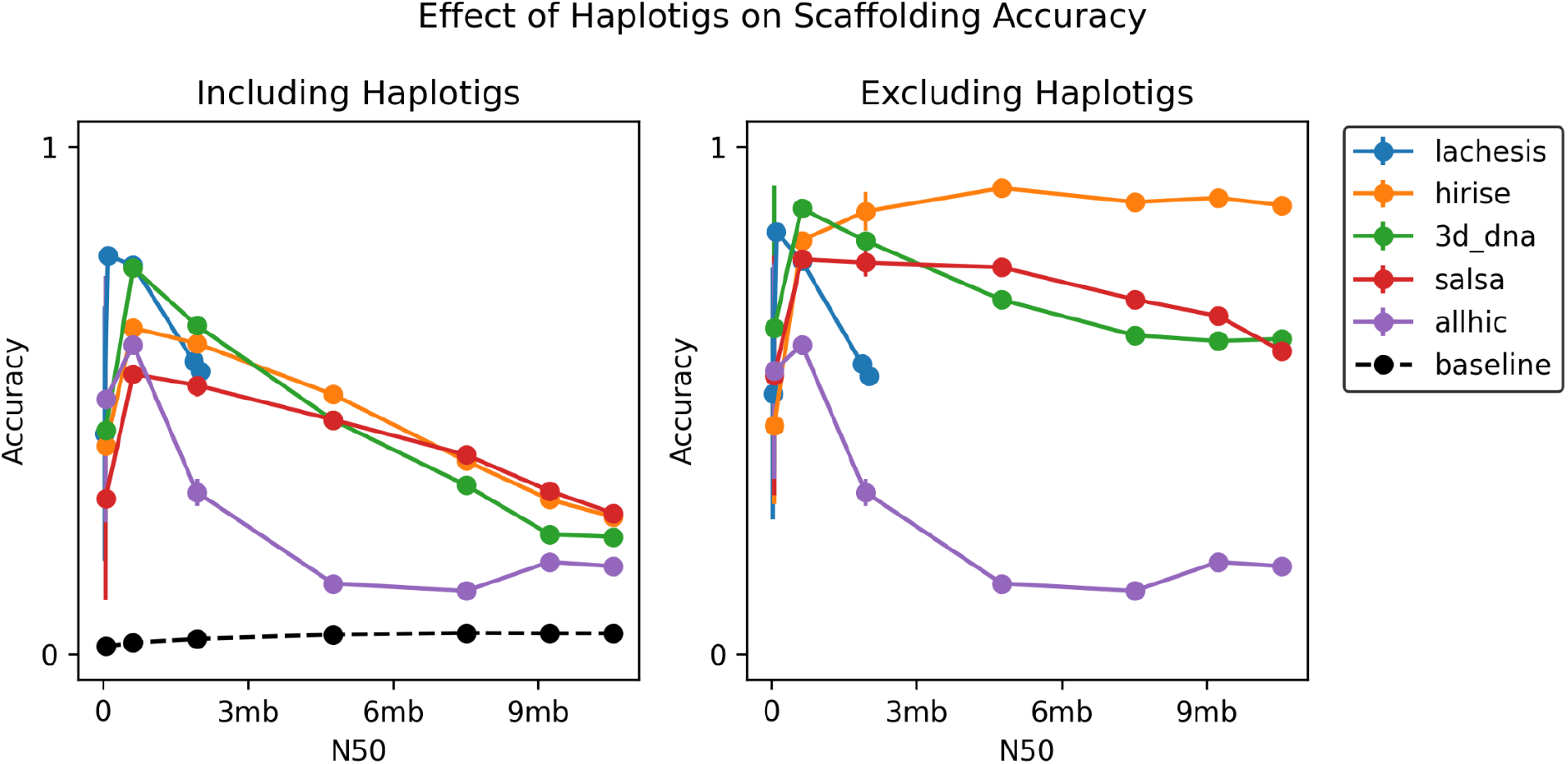
The improvement in accuracy by removing unscaffolded contigs of *H. sapiens*. We found that small overlapping contigs known as haplotigs are often excluded from scaffolds by the methods we tested. This suggests that the remaining scaffolded contigs show much higher accuracy, and is more consistent with results from the split reference setting.

While investigating the lower performance on the *de novo* assemblies of *A. thaliana* and *H. sapiens* compared to the split reference setting, we discovered that approximately fifty percent of the Hi-C reads for the *A. thaliana* assemblies mapped to more than one location, in comparison to about ten percent for the species (Supplementary Figure 14). Because multi-mapping reads cannot be reliably used in the scaffolding process, Hi-C scaffolders typically mask these regions during preprocessing. We speculate that this led to the poor scaffolding performance on *A. thaliana* assemblies (Supplementary Table 3).

While the culprit for low accuracy on *de novo A. thaliana* assemblies was the high repeat content, the same cannot be said for *H. sapiens*, which shows a multi-mapping rate on par with the other two species. Instead, our Mummer alignments of scaffolds hinted at a high frequency of small, overlapping contigs, known as haplotigs, which are typically caused by allelic variation (Supplementary Figure 11). Haplotigs are known to interfere in scaffolding since they create a scenario in which two different sequences belong to the same location along the genome. Since the goal of most assembly projects is to produce a haploid representation of the genome, haplotigs can be reasonably omitted from analysis. We found that most Hi-C scaffolders automatically exclude haplotigs in scaffolding, because they are small and therefore contribute a relatively small amount of Hi-C signal compared to the primary contig in a region. Consequently, excluding unscaffolded regions of the assembly led to only a 5% reduction in assembly size on average. The scaffolding accuracy of these assemblies improved dramatically for HiRise, Salsa, and 3d-dna, suggesting that the presence of haplotigs can obfuscate the assessment of *de novo* assemblies. As such, future studies should attempt to remove haplotypes before scaffolding. We have had preliminary success along these lines using the purge_dups pipeline (Supplementary Figure 13) [17].

## Discussion

Given the proliferation of methods to scaffold genomes using Hi-C, it is critical to understand the landscape of their performance and assess the state of the field. Overall, we found that the performance of existing Hi-C scaffolding tools varies with species and assembly size, but with two salient trends across Hi-C scaffolders. First, accuracy decreases at low N50 values. Most scaffolders join contigs by pairing the two contigs with the highest number of connecting Hi-C reads. In a context where the contigs are both small and numerous, this approach leads to ambiguities in edge strength and subsequent erroneous adjacencies. However, more surprising is the second trend, where scaffolders also perform worse in the high N50 regime. For example, when the yeast genome is broken into 1mb contigs, there are only four joins necessary to complete the genome. Yet only one assembler is successful at this task, whereas the others perform worse than the baseline of no scaffolding. Often it seems that over-grouping is the culprit, with AllHiC and 3d-dna prone to producing mega-scaffolds by grouping all of its contigs into a single scaffold. This suggests that single-contig scaffolds were not considered as a unique and important edge case for scaffolders. Recent advances in long read technologies have spurred the growth of near chromosome-length contigs in some species, and care should be taken to ensure that Hi-C scaffolders can also function as a polishing tool in this setting.

Overall, we found that HiRise offers the best performance on average across all conditions. (Table 3) Though it was the slowest of all scaffolders, it was one of the only scaffolders not to experience any significant performance decay at large N50s. It should be noted that the original software release of HiRise contains several errors in the source code, and that subsequent development has been taken on by a private company (Dovetail Genomics). Lachesis was the second best scaffolder on average, though it was the tool that most commonly failed to run under default settings. Its initial release also contains several installation bugs, and its later development has been taken on by another private company (Phase Genomics). AllHiC and Salsa yielded slightly worse performance than Lachesis. Interestingly, though 3d-dna had some of the lowest performing runs of the group under certain conditions, it remains one of the most heavily used methods. We traced its largest drop in performance to low grouping accuracy, which in turn was caused by a strong tendency to place most contigs into one or two scaffolds.

We offer several recommendations to consider when developing new Hi-C scaffolders. First, the most common starting point for Hi-C scaffolders is the BAM alignment file and assembly FASTA file. Since these files are both straightforward to parse and routinely produced, they offer excellent starting points for scaffolding. We found that deviations from this workflow, such as 3d-dna’s requirement of the Juicer pipeline, created additional barriers to use. Second, we found the requirement for chromosome count to be unnecessary to achieve good performance. The two scaffolders which required this parameter, Lachesis and AllHiC, did not perform substantially better than other methods. Because Hi-C scaffolding is most often used in a *de novo* context, chromosome count is often unknown or unreliable, and therefore should be estimated by the method itself. Third, several scaffolders include integrated methods to break scaffolds at positions where a misassembly may have occurred. This step should be optional, because in some situations it erroneously breaks contigs that have been assembled with the aid of additional sequencing information such as mate pairs or optical mapping. Finally, scaffolders should output an AGP file to describe the organization of contigs [18]. Several scaffolders omit this information, and some create their own bespoke file formats which lead to problems comparing and understanding scaffolding outputs.

The current state of Hi-C scaffolding remains a two-step process: first, assemblies are passed through scaffolding software, and then the errors are fixed by hand. Opportunities for improvement lie on both ends of the workflow - better algorithms to scaffold genomes with Hi-C, and more modern tools for the manual correction of scaffolds. Despite any shortcomings of current methods, Hi-C scaffolding is a powerful tool in the evolving science of building reference genomes, and we hope that future developers can look to our study to help select an appropriate scaffolder for their own assembly tasks.

## Acknowledgements

We thank Shawn Sullivan of Phase Genomics for his extensive advice and support. Working with Hi-C data, and assemblies in particular remains somewhat of an art form, and Shawn’s experience with hundreds of libraries and species proved invaluable. Funding for A.S. was obtained in part from 6R01AI103858-06 from NIAID and Seed funding from Seattle Children’s Research Institute to P. M.. This work was facilitated through the use of advanced computational, storage, and networking infrastructure provided by the Hyak supercomputer system and funded by the STF at the University of Washington.

## References

[1] Burton, Joshua N., et al. “Chromosome-scale scaffolding of de novo genome assemblies based on chromatin interactions.” Nature biotechnology 31.12 (2013): 1119–1125.

[2] Kaplan, Noam, and Job Dekker. “High-throughput genome scaffolding from in vivo DNA interaction frequency.” Nature biotechnology 31.12 (2013): 1143–1147.

[3] Marie-Nelly, Hervé, et al. “High-quality genome (re) assembly using chromosomal contact data.” Nature communications 5.1 (2014): 1–10.

[4] Putnam, Nicholas H., et al. “Chromosome-scale shotgun assembly using an in vitro method for long-range linkage.” Genome research 26.3 (2016): 342–350.

[5] Dudchenko, Olga, et al. “De novo assembly of the Aedes aegypti genome using Hi-C yields chromosome-length scaffolds.” Science 356.6333 (2017): 92–95.

[6] Ghurye, Jay, et al. “Integrating Hi-C links with assembly graphs for chromosome-scale assembly.” PLoS computational biology 15.8 (2019): e1007273.

[7] Zhang, Xingtan, et al. “Assembly of allele-aware, chromosomal-scale autopolyploid genomes based on Hi-C data.” Nature plants 5.8 (2019): 833–845.

[8] Renschler, Gina, et al. “Hi-C guided assemblies reveal conserved regulatory topologies on X and autosomes despite extensive genome shuffling.” Genes & development 33.21-22 (2019): 1591–1612.

[9] Nakabayashi, Ryo, and Shinichi Morishita. “HiC-Hiker: a probabilistic model to determine contig orientation in chromosome-length scaffolds with Hi-C.” Bioinformatics 36.13 (2020): 3966–3974.

[10] Koren, Sergey, et al. “Canu: scalable and accurate long-read assembly via adaptive k-mer weighting and repeat separation.” Genome research 27.5 (2017): 722–736.

[11] Li, Heng. “Aligning sequence reads, clone sequences and assembly contigs with BWA-MEM.” arXiv preprint arXiv:1303.3997 (2013).

[12] Faust, Gregory G., and Ira M. Hall. “SAMBLASTER: fast duplicate marking and structural variant read extraction.” Bioinformatics 30.17 (2014): 2503–2505.

[13] Danecek, Petr, et al. “Twelve years of SAMtools and BCFtools.” Gigascience 10.2 (2021): giab008.

[14] Durand, Neva C., et al. “Juicer provides a one-click system for analyzing loop-resolution Hi-C experiments.” Cell systems 3.1 (2016): 95–98.

[15] Marçais, Guillaume, et al. “MUMmer4: A fast and versatile genome alignment system.” PLoS computational biology 14.1 (2018): e1005944.

[16] Bergeron, Anne, Julia Mixtacki, and Jens Stoye. “A unifying view of genome rearrangements.” International Workshop on Algorithms in Bioinformatics. Springer, Berlin, Heidelberg, 2006.

[17] Guan, Dengfeng, et al. “Identifying and removing haplotypic duplication in primary genome assemblies.” Bioinformatics 36.9 (2020): 2896–2898.

[18] NCBI. Agp specification v2.1. National Center for Biotechnology Information, 2019.

