## Supplementary Figures for "A benchmark of Hi-C scaffolders using reference genomes and *de novo* assemblies"

### Supplement

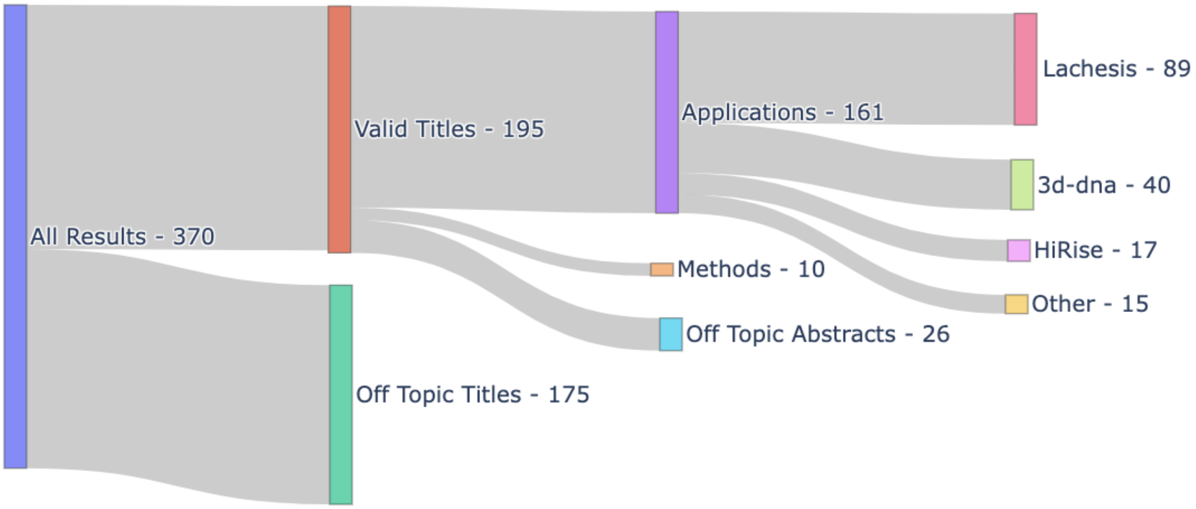


**Supplementary Table 1.** A sankey diagram depicting the literature search process. We identified ten different Hi-C scaffolding methods in the literature and found that their usage varied significantly, with only five methods showcasing more than three published genomes.

| | Grouping | | Order | | Orientation | | Accuracy | |
|---|---|---|---|---|---|---|---|---|
| **Method** | **Split** | **De novo** | **Split** | **De novo** | **Split** | **De novo** | **Split** | **De novo** |
| lachesis | **0.91 ± 0.1** | **0.71 ± 0.2** | **0.95 ± 0.03** | **0.66 ± 0.1** | **0.95 ± 0.03** | **0.55 ± 0.2** | **0.95 ± 0.03** | 0.55 ± 0.2 |
| hirise | 0.74 ± 0.3 | 0.69 ± 0.2 | 0.87 ± 0.2 | 0.49 ± 0.3 | 0.87 ± 0.2 | 0.44 ± 0.3 | 0.87 ± 0.2 | **0.62 ± 0.2** |
| 3d_dna | 0.35 ± 0.2 | 0.38 ± 0.3 | 0.66 ± 0.3 | 0.34 ± 0.2 | 0.66 ± 0.3 | 0.30 ± 0.2 | 0.71 ± 0.2 | 0.41 ± 0.2 |
| salsa | 0.52 ± 0.3 | 0.47 ± 0.3 | 0.85 ± 0.2 | 0.44 ± 0.2 | 0.84 ± 0.2 | 0.41 ± 0.2 | 0.85 ± 0.1 | 0.50 ± 0.2 |
| allhic | 0.75 ± 0.4 | 0.52 ± 0.3 | 0.95 ± 0.1 | 0.58 ± 0.3 | 0.94 ± 0.1 | 0.49 ± 0.3 | 0.84 ± 0.2 | 0.39 ± 0.3 |
| baseline | 0.18 ± 0.3 | 0.36 ± 0.3 | 0.03 ± 0.1 | 0 | 0.03 ± 0.1 | 0 | 0.16 ± 0.2 | 0.28 ± 0.3 |

**Supplementary Table 2.** An overview of performance of each of the methods based on their average accuracy determined by Edison. The split column refers to the task of scaffolding equal sized pieces of the reference genome. The *de novo* column refers to the task of scaffolding the assemblies created by Canu. Baseline represents the score for contigs without scaffolding.

| Organism | Genome Size | Reads | Bases | Coverage | BioProject |
|---|---|---|---|---|---|
| S. cerevisiae | 12,100,000 | 313,114 | 1,701,530,052 | 141 | PRJEB7245 |
| L. tarentolae | 32,200,000 | 1,360,815 | 7,198,339,498 | 224 | PRJNA821548 |
| A. thaliana | 135,000,000 | 7,353,356 | 49,942,606,909 | 370 | PRJNA314706 |
| H. sapiens | 3,100,000,000 | 47,885,330 | 328,978,598,683 | 106 | PRJNA301527 |

**Supplementary Table 3.** Overview of data collected for *de novo* genome assemblies. The amount of data is roughly proportional to the size of the genome such that they can be down-sampled to a similar read coverage.

| Coverage | Yeast N50 | Leishmania N50 | Arabidopsis N50 | Human N50 |
|---|---|---|---|---|
| 10 | 21,825 | 9,250 | 24,823 | 18,711 |
| 20 | 176,516 | 18,595 | 86,731 | 43,231 |
| 30 | 551,752 | 41,537 | 159,879 | 105,563 |
| 40 | 568,123 | 40,610 | 147,845 | 626,886 |
| 50 | 614,056 | 94,577 | 149,418 | 1,873,143 |
| 60 | 813,309 | 213,275 | 161,138 | 2,018,914 |
| 70 | 777,771 | 71,335 | 140,209 | 4,761,131 |
| 80 | 813,629 | 117,224 | 149,909 | 7,508,518 |
| 90 | 813,427 | 229,090 | 164,531 | 9,231,632 |
| 100 | 930,538 | 332,478 | 186,445 | 10,553,285 |

**Supplementary Table 4.** Overview of the *de novo* assemblies created by Canu. We generated ten assemblies for each species and downsampled reads used to create them to vary their N50s. As a general trend, we observed that increased read coverage led to long contigs. Arabidopsis appeared to be an outlier of this trend, and its genome assembly appeared to be challenging and indicative of high rates of heterozygosity.

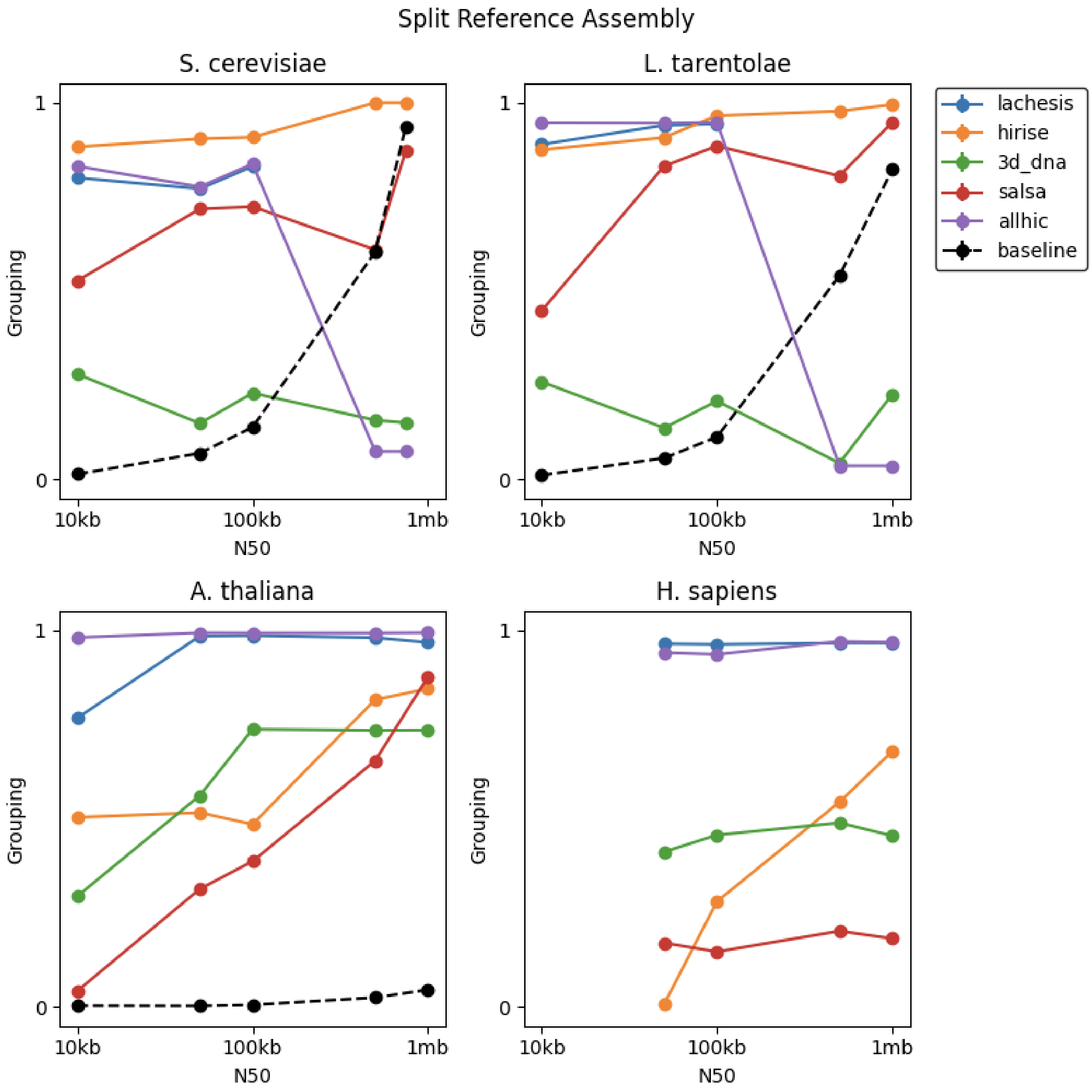


**Supplement 4.** The grouping scores of scaffolders on split reference assemblies. There is wide variation in grouping performance, with trends pointing to difficulty with small assemblies with large N50s and large assemblies with small N50s.

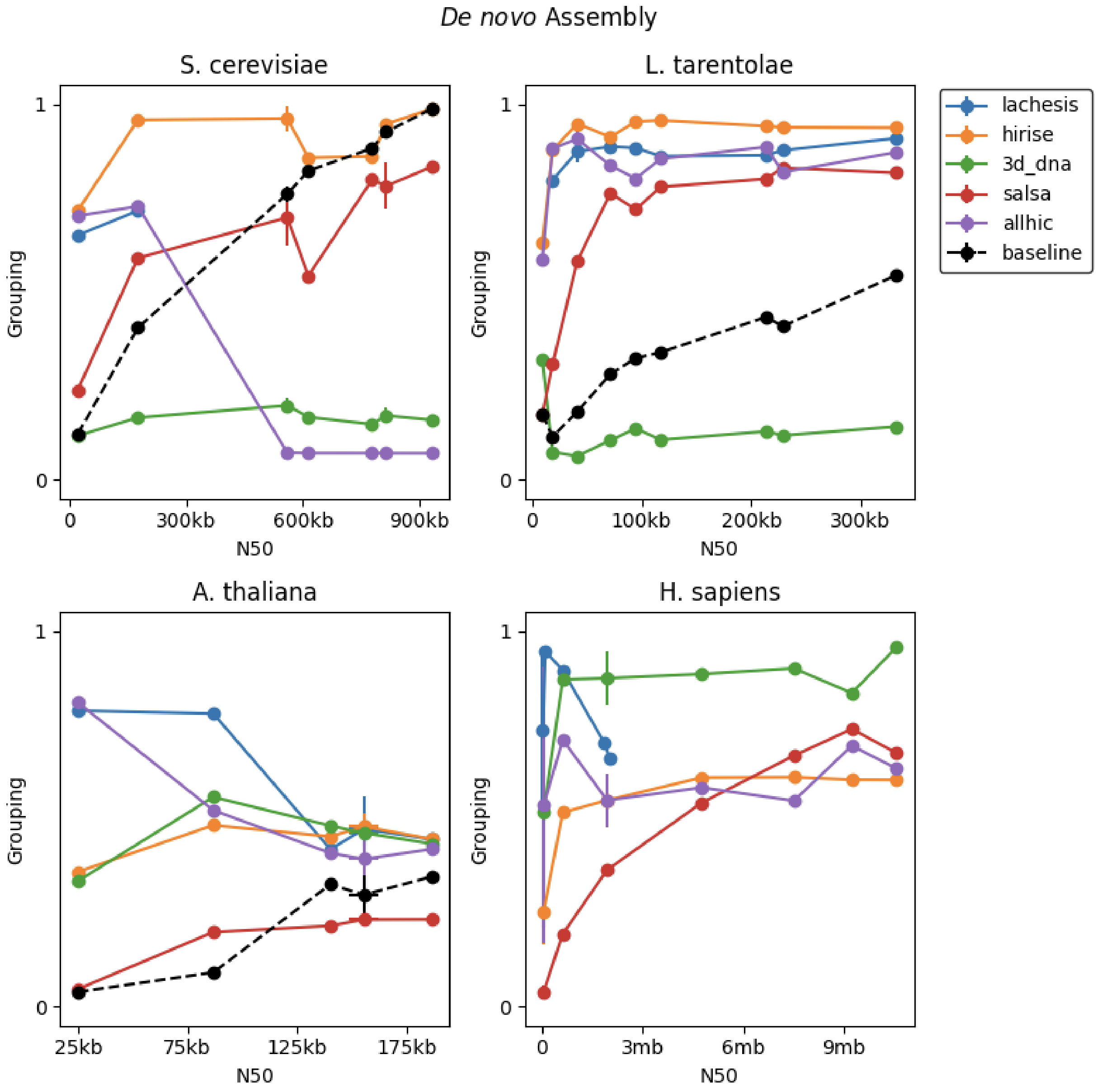


**Supplement 5.** The grouping scores for Hi-C scaffolders on *de novo* assemblies. Higher grouping accuracy indicates that scaffolders were able to uniquely isolate contigs belonging to the same chromosome within scaffolds.

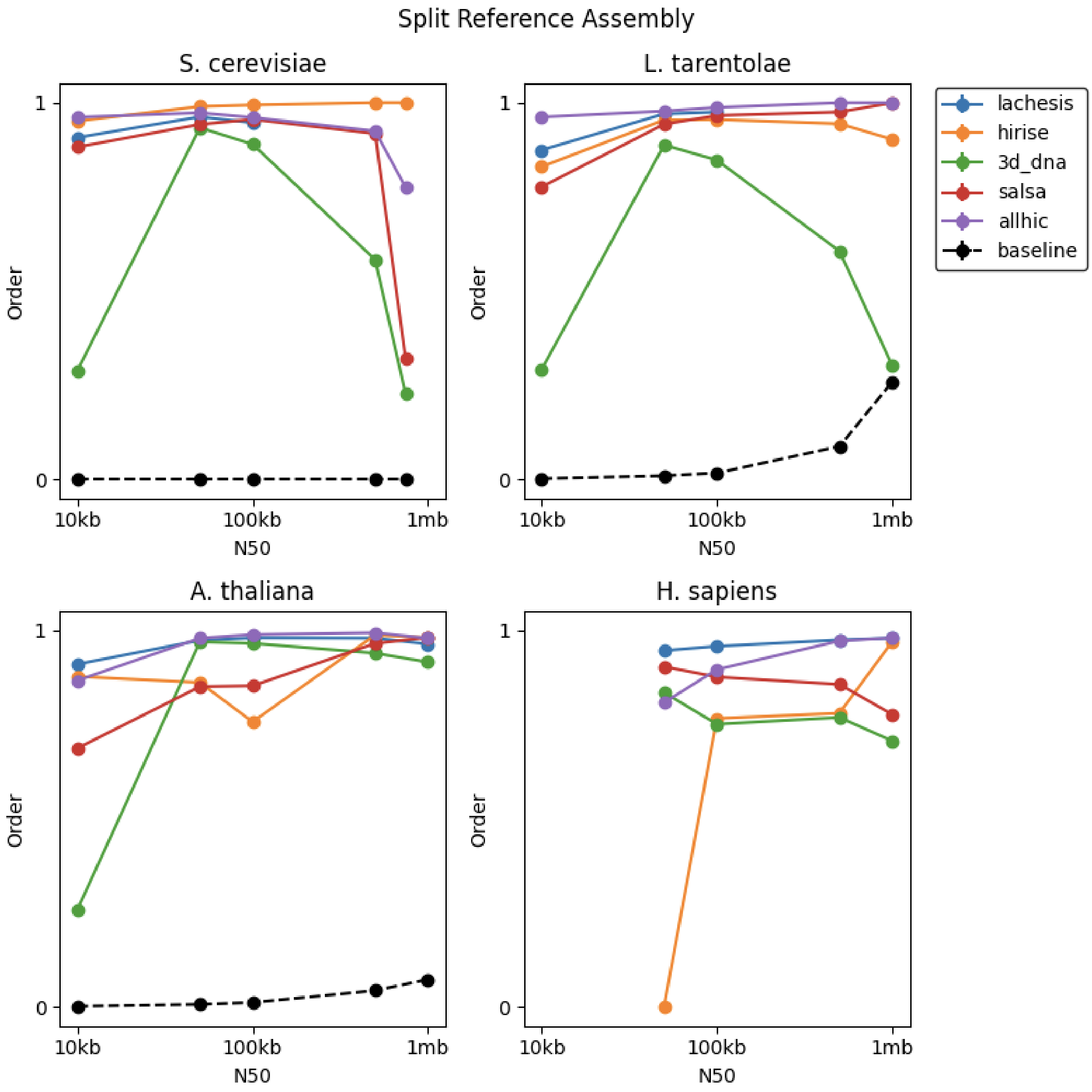


**Supplement 6**. The order scores for Hi-C scaffolders on split reference assemblies.

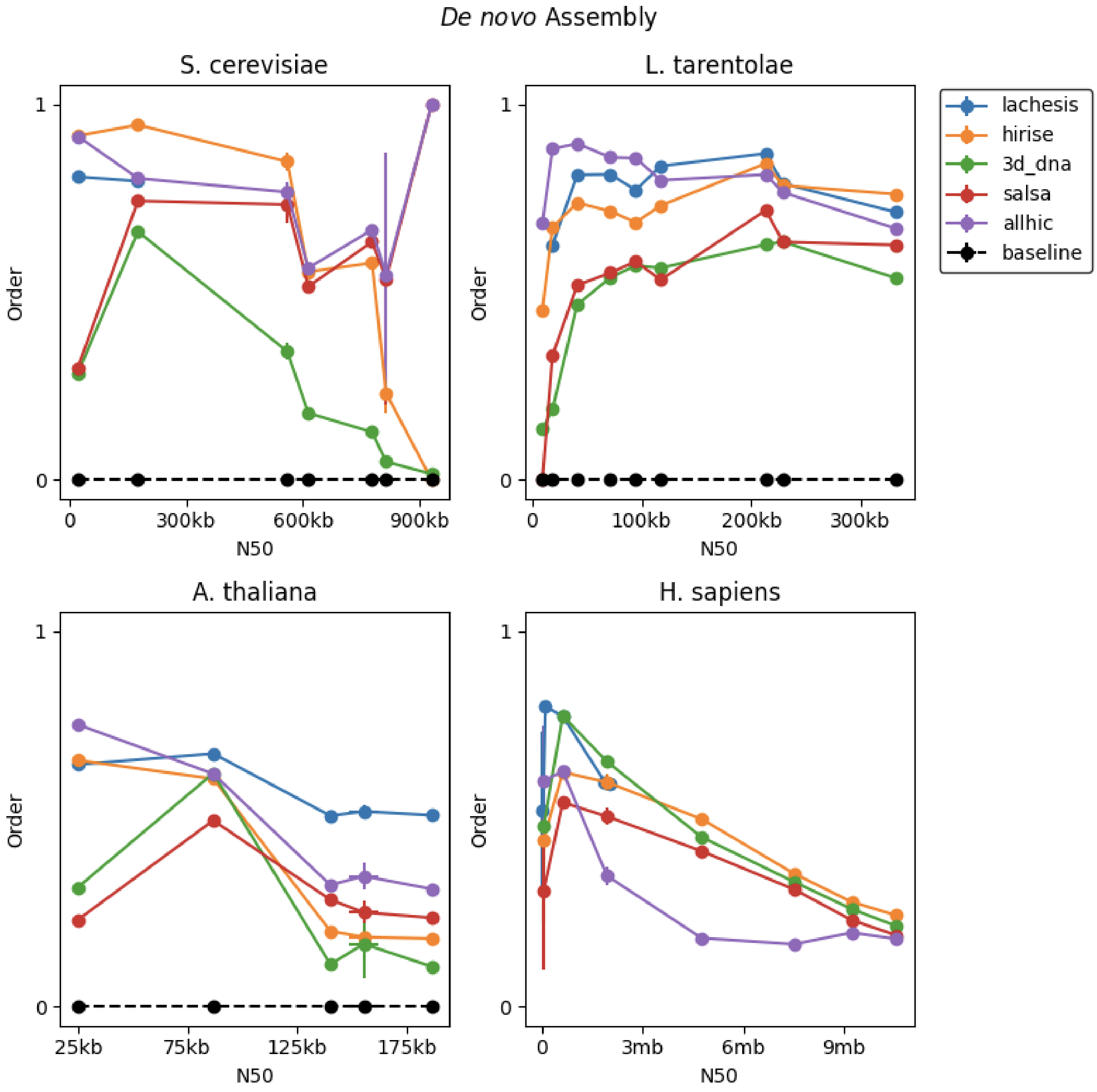


**Supplement 7.** The order scores for Hi-C scaffolders on *de novo* assemblies. Higher order accuracy indicates that scaffolders were able to correctly place contigs next to their expected neighbors.

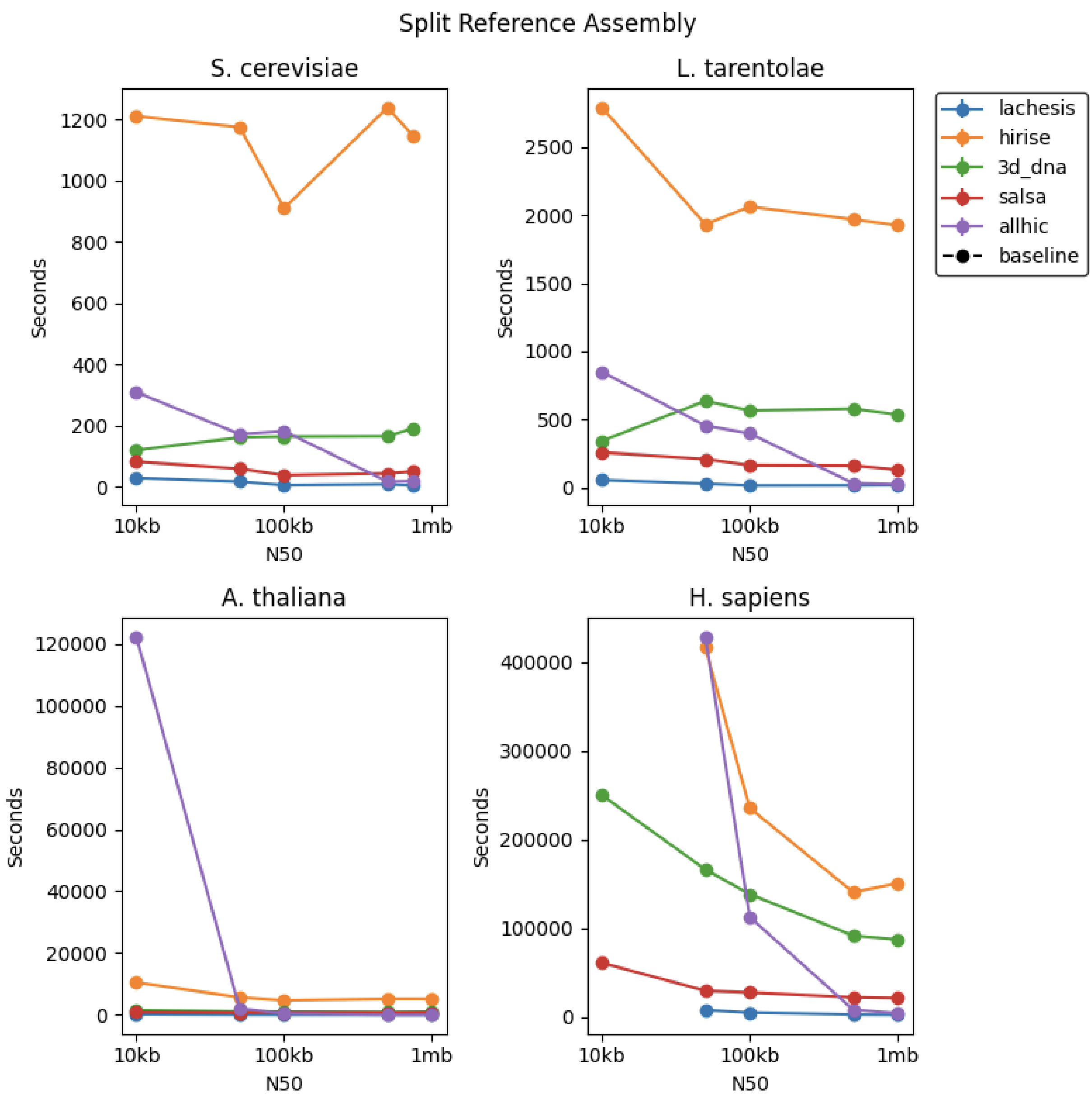


**Supplement 8**. The runtime of Hi-C scaffolders on split assemblies.

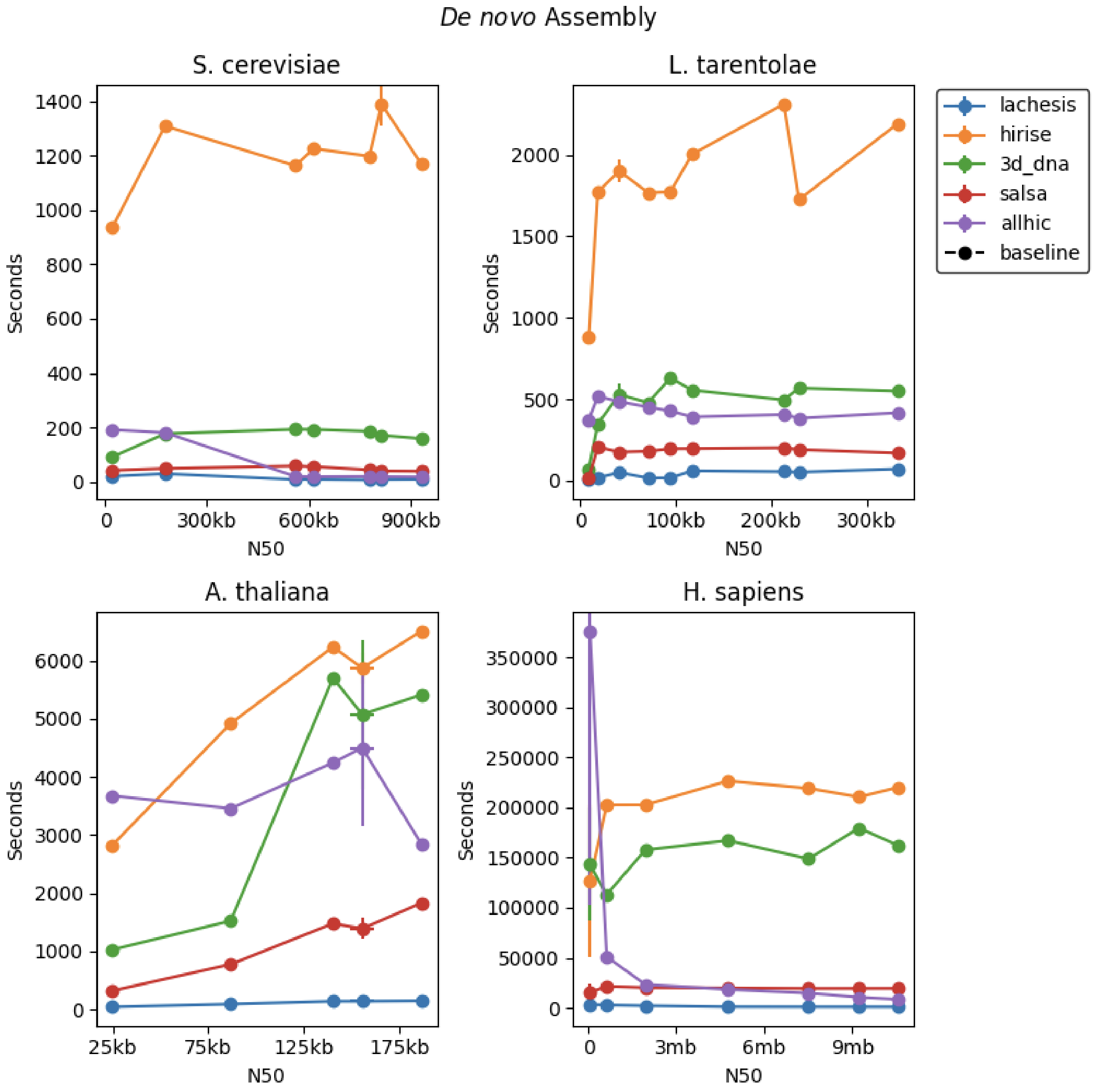


**Supplement 9.** The runtime of Hi-C scaffolders on *de novo* assemblies. Hirise is generally the slowest and Lachesis the fastest scaffolder.

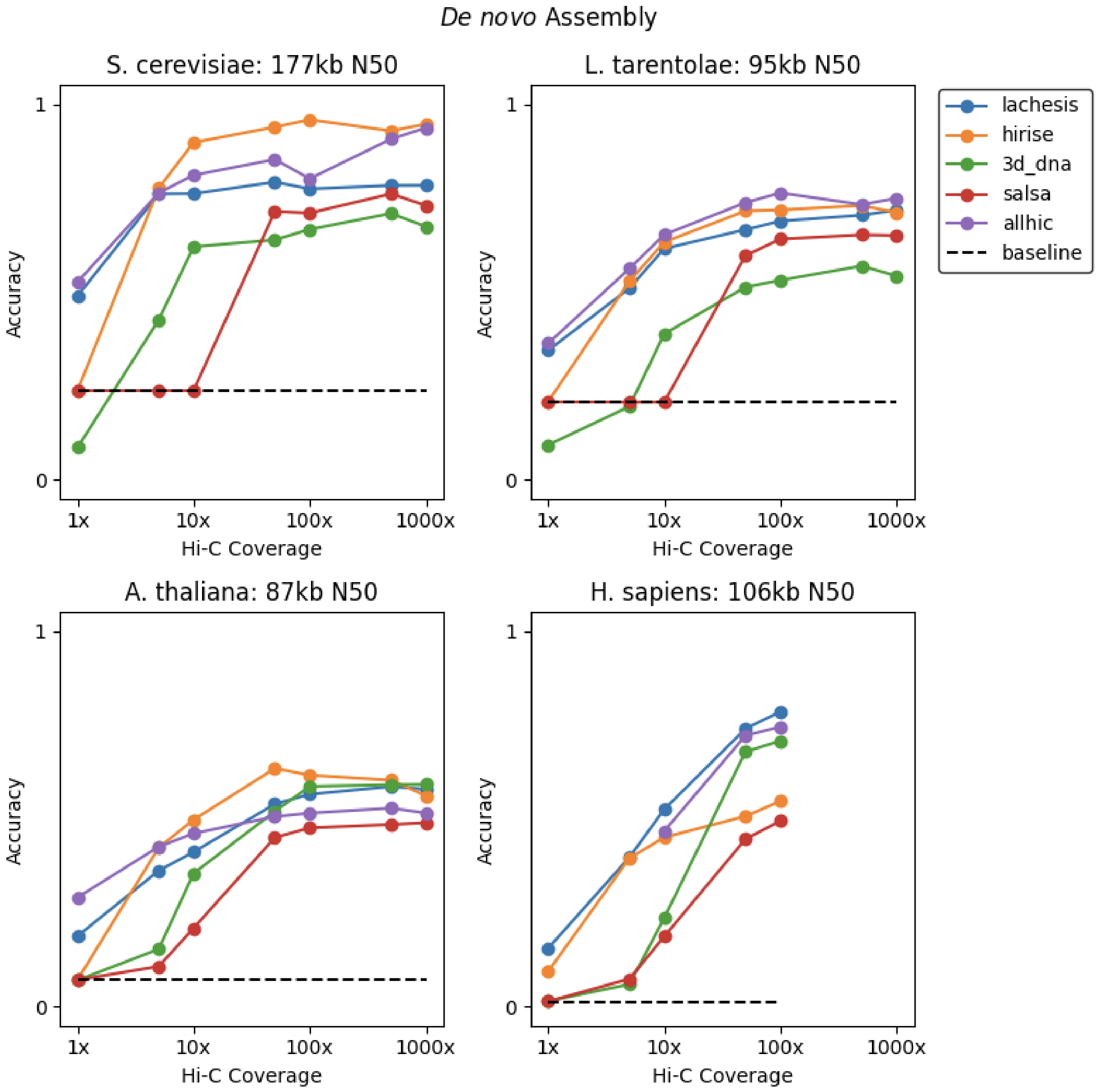


**Supplement 10.** Downsampling of Hi-C reads on *de novo* assemblies. The same trend as the split references is seen here, where Hi-C read densities below 50 reads per kilobase lead to a decline in performance.

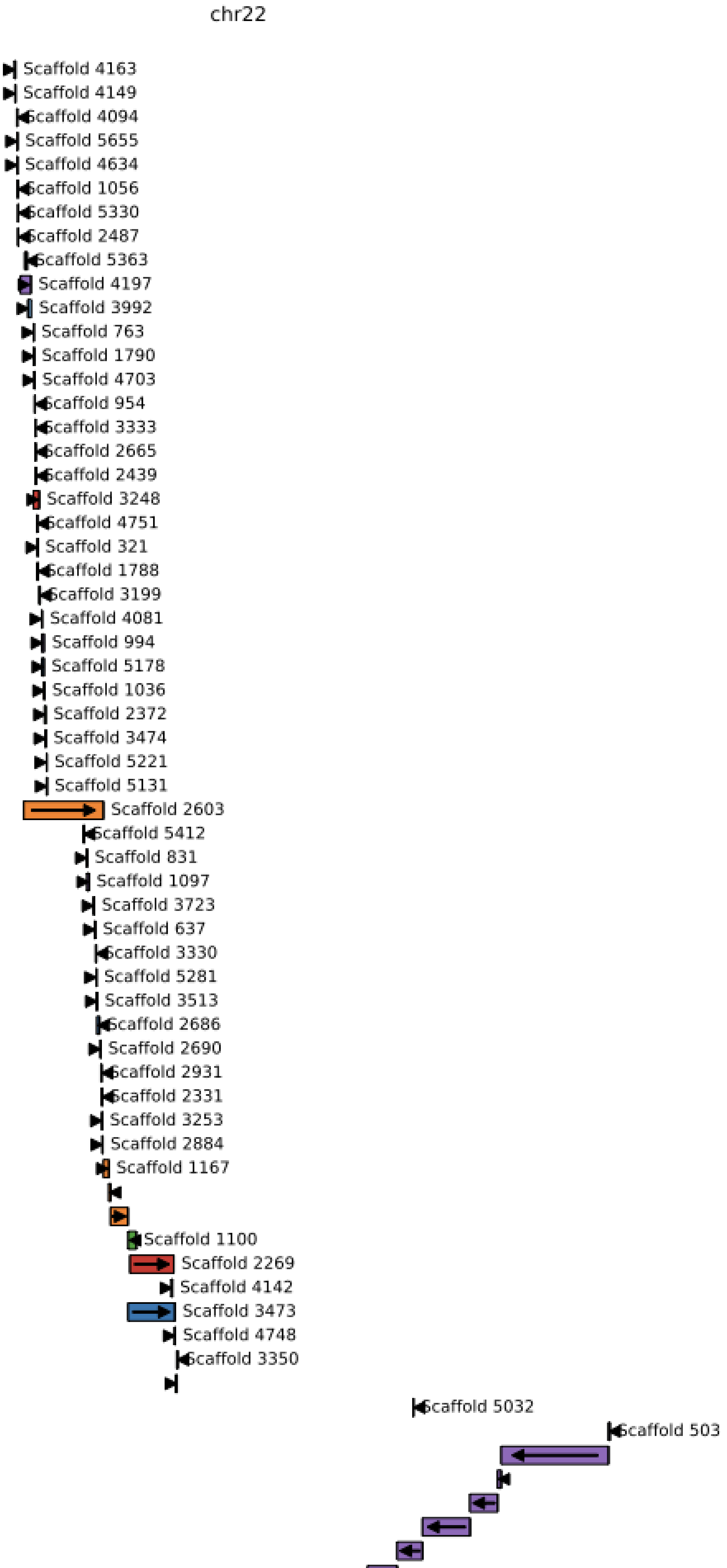

chr22
Scaffold 4163
Scaffold 4149
Scaffold 4094
Scaffold 5655
Scaffold 4634
Scaffold 1056
Scaffold 5330
Scaffold 2487
Scaffold 5363
Scaffold 4197
Scaffold 3992
Scaffold 763
Scaffold 1790
Scaffold 4703
Scaffold 954
Scaffold 3333
Scaffold 2665
Scaffold 2439
Scaffold 3248
Scaffold 4751
Scaffold 321
Scaffold 1788
Scaffold 3199
Scaffold 4081
Scaffold 994
Scaffold 5178
Scaffold 1036
Scaffold 2372
Scaffold 3474
Scaffold 5221
Scaffold 5131
Scaffold 2603
Scaffold 5412
Scaffold 831
Scaffold 1097
Scaffold 3723
Scaffold 637
Scaffold 3330
Scaffold 5281
Scaffold 3513
Scaffold 2686
Scaffold 2690
Scaffold 2931
Scaffold 2331
Scaffold 3253
Scaffold 2884
Scaffold 1167
Scaffold 1100
Scaffold 2269
Scaffold 4142
Scaffold 3473
Scaffold 4748
Scaffold 3350
Scaffold 5032
Scaffold 503

**Supplement 11.** An overview of how HiRise scaffolded the 10mb N50 *H. sapiens* assembly for Chromosome 22. Each row represents a contig, and each label and color represents a scaffold. The x-axis represents the alignment based position of the contig, and the y-axis represents the scaffolder based order of the contigs. Here, HiRise picks out a set of larger contigs and scaffolds them in the correct order relative to each other. However it leaves out a number of the smaller contigs, including ones that overlap with its primary scaffold (Scaffold 503) for this chromosome.

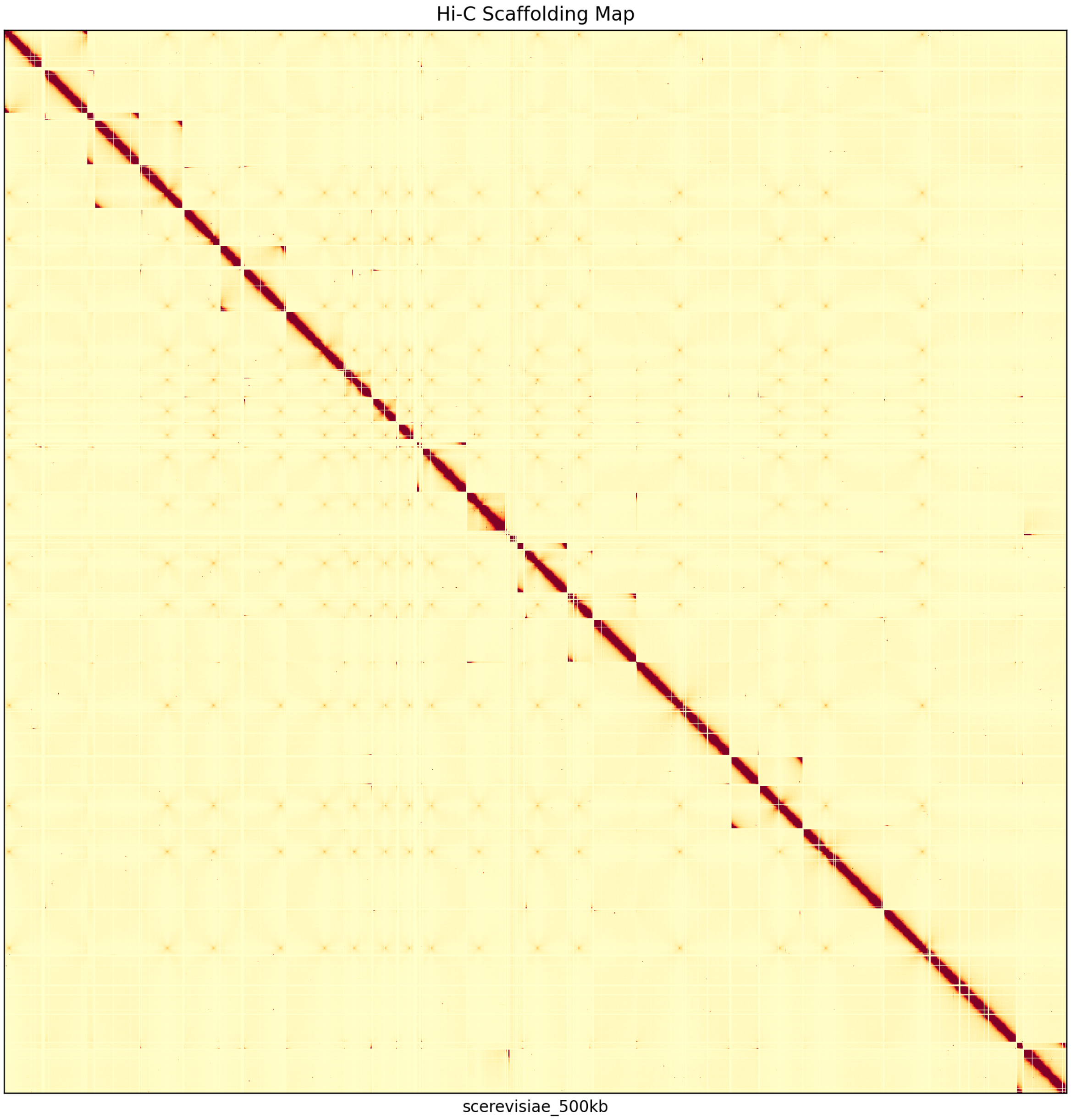


**Supplement 12.** AllHiC scaffolding 500kb contigs from the split reference assembly of *S. cerevisiae.* While the contigs have been placed mostly in the correct order and orientation, all the contigs were placed in a single mega-scaffold causing the overall accuracy to dramatically decrease for this particular scaffolding.

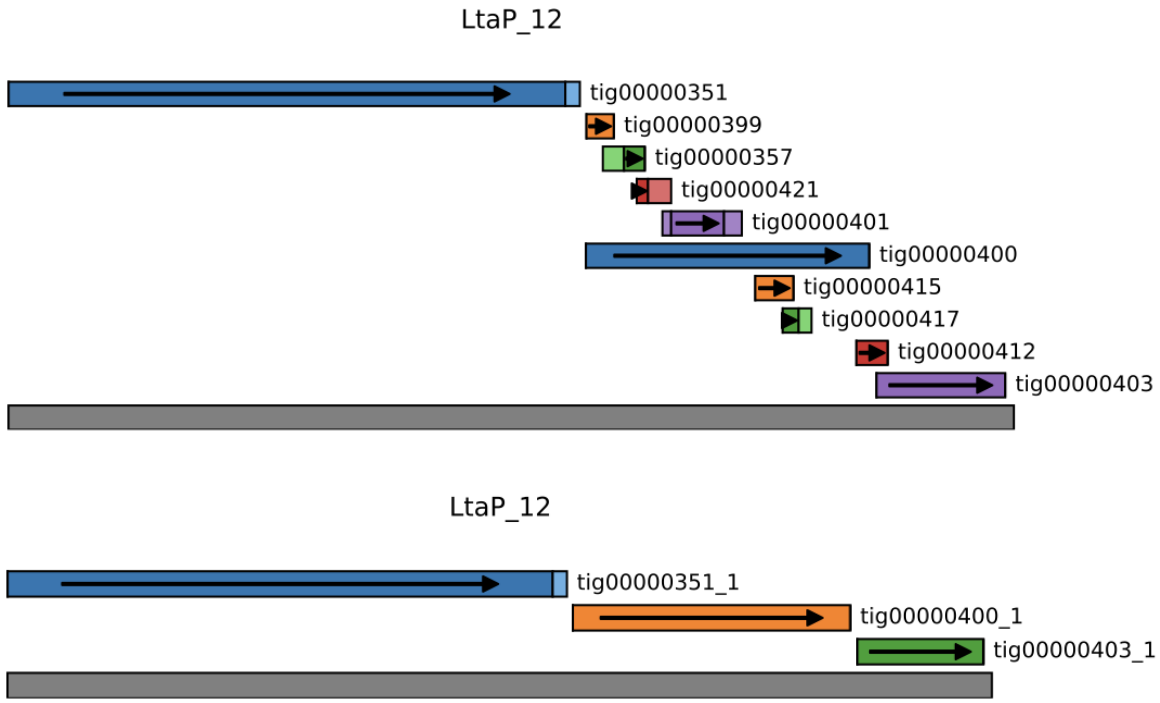


**Supplementary Figure 13.** Using purge_dups to remove halpotigs. The top section shows the contigs of the original assembly for L. tarentolae that map to chromosome 12. The bottom section shows the remaining contigs after the purging of haplotigs.

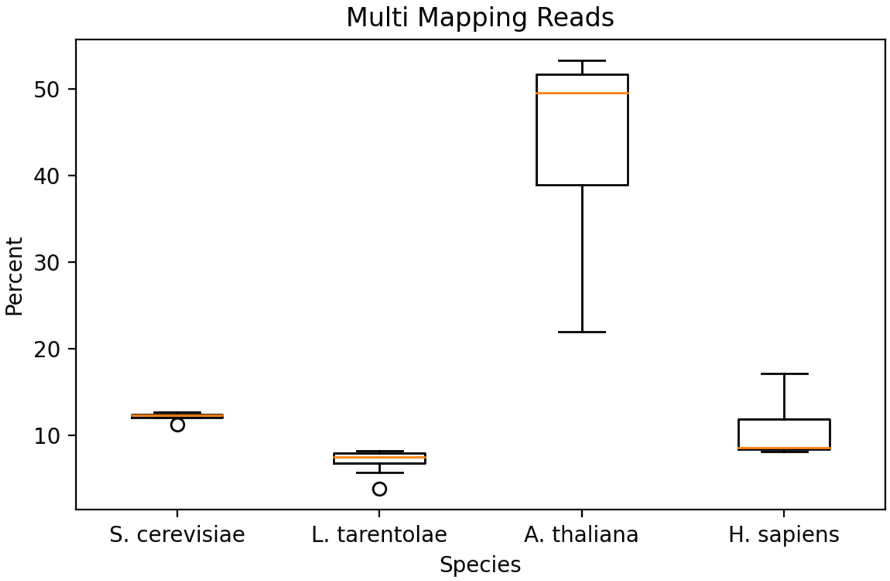


**Supplementary Figure 14.** The percent of reads that map to multiple positions in the *de novo* assembly. We found that as the number of reads used to create the *de novo* assembly goes up, the repetitive content of the genome goes up. The uniformly low accuracy against *A. thaliana* assemblies can likely be attributed to a high percentage of multi-mapping reads, which cannot be used by Hi-C scaffolders.
